# Candidate genes for stem rust resistance in Italian ryegrass revealed by nested association mapping

**DOI:** 10.64898/2025.12.19.695465

**Authors:** Jenny Kiesbauer, Christoph Grieder, Daniel Ariza-Suarez, Maria Hug, Linda Helene Schlatter, Meril Sindelar, Dario Copetti, Bruno Studer, Roland Kölliker

## Abstract

Stem rust, caused by *Puccinia graminis* ssp. *graminicola*, is a major disease affecting the outcrossing species Italian ryegrass (*Lolium multiflorum* Lam.), leading to substantial reductions in seed yield. Until now, knowledge on the genetic control of stem rust resistance in Italian ryegrass has been limited to a few quantitative trait loci identified in bi-parental mapping populations. To discover novel resistance sources for use in breeding programs, appropriate plant populations, reliable phenotyping methods and advanced genomic tools are essential.

In this study, we utilized a previously established F_2_ nested association mapping (NAM) population comprising 708 individuals derived from 24 founder plants exhibiting high variation in stem rust resistance. Phenotypic evaluation was conducted under natural inoculation in three location-by-year combinations. By integrating reduced-representation sequencing of the NAM population with whole-genome sequencing of the founder plants, we identified 3,199,253 SNP markers for association mapping. The high SNP marker density, together with the strong detection power of the NAM population, enabled the identification of four novel candidate genes. Two of these genes, located on chromosomes 6 and 7, encode receptor-like serine/threonine kinases that are known to play a role in stem rust resistance in other crops. Within the serine/threonine kinase gene *Chr7.32208*, two superior haplotypes were identified that can be directly implemented as selection criteria in breeding programs. The novel stem rust resistance candidate genes reported here provide promising targets for functional validation and the improvement of stem rust resistance in Italian ryegrass breeding.

**Key message:** Field phenotyping of a previously established NAM population for stem rust resistance, combined with high-density genotyping, enabled the identification of novel sources of stem rust resistance in Italian ryegrass.

## Introduction

Italian ryegrass (*Lolium multiflorum* Lam.) is one of the main constituents of productive grassland systems, valued for its high biomass yield potential, fast ground cover and excellent digestibility for ruminants. Breeding of Italian ryegrass relies on population breeding with a main focus on forage-related traits like biomass yield, digestibility, palatability and disease resistance (Lüscher et al. 2019). In addition, traits related to seed yield are important for seed multiplication of cultivars. For seed yield and related traits, resistance to stem rust is of particular importance, as this disease can reduce seed yield by up to 35% (Hampton 1986; Pfender 2009; Schubiger et al. 2010).

Stem rust is caused by *Puccinia graminis* ssp. *graminicola*, an obligate biotrophic fungus that infects stems and leaf sheaths of susceptible ryegrass plants at seed formation (Leonard and Szabo 2005). One option to reduce stem rust and, therefore, ensure a high seed yield in seed production, is the use of fungicides which can reduce but not entirely prevent infections (Rodriguez-Algaba et al. 2020). A more sustainable method of preventing stem rust infections is to breed for disease resistant cultivars (Schubiger et al. 2010). Higher selection efficiency and precision can be achieved by marker-assisted selection (Collard et al. 2005). For example, a single marker closely linked to crown rust resistance caused by *P. coronata* f. sp. *lolii* was successfully used to improve crown rust resistance within breeding germplasm of Italian ryegrass (Kölliker et al. 2016). To prevent recombination from disrupting the association between a single marker and its corresponding causal gene or regulatory element, the marker needs to be physically close to the causal locus. Haplotypes are defined by combinations of multiple markers within a genomic region of the same chromosome segment which tend to inherit together (Bernardo 2010). Compared to single markers, haplotypes capture a broader spectrum of genetic variation and, therefore, enhance the resolution of potential genomic areas and provide a more effective and stable tool for identifying superior allelic combinations (Stephens et al. 2001).

In the last decades, several loci associated with stem rust resistance have been identified in ryegrass. To date, quantitative trait loci (QTL) conferring resistance against stem rust were found on linkage group (LG) 4 using an interspecific (Italian ryegrass x perennial ryegrass) cross (Jo et al. 2008). In addition, three pathotype-specific resistance QTL were found on LG 1, 6 and 7 in a perennial ryegrass population (Pfender et al. 2011; Pfender and Slabaugh 2013). Furthermore, a qualitative resistance locus on LG 4, named *LpPg1*, was found in perennial ryegrass (Beckmann 2010). Bulk expression studies using the same biparental mapping population as Beckmann (2010) revealed a NBS-LRR candidate resistance gene co-segregating with the *LpPg1* resistance locus (Bojahr et al. 2016). All the previous QTL mapping studies were conducted based on F_1_ biparental populations, using one parent as resistance and the other one as susceptibility source. In general, QTL mapping based on F_1_ biparental populations within self-incompatible outcrossing species has a high statistical power to examine loci associated with the trait within the specific population and was, therefore, successfully implemented in ryegrasses for metabolites (Foito et al. 2017), vernalization response (Jensen et al. 2005) and disease resistance (Studer et al. 2006, 2007). The transfer of QTL based on biparental populations into breeding programs may be difficult due to different genetic backgrounds and unfavorable alleles present in the mapping populations. Consequently, such QTL are rarely used in the development of resistant cultivars. To find causal genes rather than loci associated with disease resistance, suitable plant populations, advanced genomic tools and a reliable phenotyping method are prerequisites.

Nested association mapping (NAM) populations combine the advantages of high statistical power and high mapping resolution (Yu et al. 2008). Such NAM populations are designed by crossing multiple diverse parents with one common founder parent. Compared to a genome wide association mapping (GWAS) population, a NAM population has the advantage of a higher detection power while maintaining allelic diversity and a low population structure, which is an excellent prerequisite to discover loci influencing quantitative traits. In addition, the parental genotypes used for establishment of the population can be selected from breeding germplasm, making the mapping results readily transferable into breeding programs. Within this project, we took advantage of a previously established NAM population with high phenotypic variation for stem rust resistance and seed production (Kiesbauer et al. 2025). With the reduced costs for sequencing combined with the increasing availability of high-quality genome assemblies, the second prerequisite of the availability of genomic tools to identify causal resistance genes is fulfilled. Within the last years, especially in ryegrasses, several reference genomes have become available (Byrne et al. 2015; Copetti et al. 2021; Frei et al. 2021; Chen et al. 2024, 2025). For cost-efficient genotyping, various tools are now available. For example, single nucleotide polymorphisms (SNPs) derived from double digest restriction site associated DNA sequencing (ddRAD; Peterson et al. 2012) were successfully reported to identifying QTL or genes related to a trait (Scheben et al. 2020; Guden et al. 2023). A reliable and easy phenotyping method for stem rust is the third prerequisite for successful resistance breeding. Until now, stem rust phenotyping was either conducted by artificial inoculation under greenhouse conditions on single plants (Pfender et al. 2011; Pfender and Slabaugh 2013) or by using an *in vitro* leaf segment test (LST; Beckmann 2010; Bojahr et al. 2016). In comparison to artificial conditions in the greenhouse or field trials, the LST is an easier and less resource-demanding (space, equipment) approach to phenotype rust in an artificial environment (Lellbach 1994). To date, no resistance data under field conditions from the same population were systematically assessed and compared to LST data. However, a high correlation between LST and field data is necessary to use LST instead of field data for resistance breeding (Studer et al. 2007).

The overall aim of this study was to elucidate the genetic control of stem rust resistance in Italian ryegrass and to provide the basis for efficient resistance breeding. In particular, our goals were i) to evaluate a previously established NAM population for haplotypes with higher stem rust resistance, ii) to identify candidate genes associated with stem rust resistance and iii) to test whether LST results are good predictors for resistance observed in the field.

## Materials and methods

### 2.1 Plant material

The NAM population consisted of 708 plants that were generated by crossing 23 diverse founder plants to one common founder plant. As the common founder, a genotype of the cv. ‘Rabiosa’, for which a phased diploid genome assembly is available (Chen et al. 2025), was used. The 23 diverse founder genotypes were derived from five ecotype populations, eight registered cultivars and ten accessions from advanced breeding germplasm (Kiesbauer et al. 2025). From each of the 23 F_1_ families, 18 plants were randomly chosen for open pollination. From the resulting 414 F_1_ plants, two seeds each were grown to establish the NAM population. In total, 120 of the plants were lost during cultivation, resulting in a final population of 708 plants (Kiesbauer et al. 2025).

### 2.2 Phenotypic evaluation under natural infection in three trials

Phenotypic data of the NAM population was collected from field trials in Zurich (47.427 °N, 8.516 °E) in 2020 (ZH20) and 2021 (ZH21) and in Oensingen (47.284 °N, 7.730 °E) in 2021 (OE21). In each trial, two clonal replicates of each of the 708 NAM genotypes and the 24 founder genotypes were planted using an alpha design (Piepho et al. 2006). Trials were planted in late summer of the preceding season and the first growth after overwintering was cut shortly after the end of heading. Plants were fixed to bamboo sticks and grown until seeds were fully ripened. Heading date and start of flowering were determined, and the occurrence of stem rust on each plant was scored at seed harvest using a scale from 1 to 9, defined as: 1 = no rust disease, 2 = traces of rust, 3 = 5%, 4 = 10%, 5 = 25%, 6 = 40%, 7 = 60%, 8 = 75% and 9 = more than 75% of the plant covered with rust (Schubiger and Boller 2016). The seed harvesting date was determined based on the cumulative average daily temperature after the start of flowering determined from the earliest flowering plant in each trial. Once a plant reached a temperature sum of 618 °C, 598 °C and 555 °C for the trials ZH20, ZH21 and OE21, respectively, seeds were harvested.

### 2.3 Phenotypic evaluation of resistance to stem rust using a leaf segment test

#### 2.3.1 Collection and propagation of urediniospores

Urediniospores of *P. graminis* ssp. *graminicola* were collected from stem rust infected plants of the field trial ZH20 using a dustbuster (16.2 Wh Dustbuster Lithium, Black & Decker, Towson, Maryland, USA). The spores were dried for one week in a desiccator over silica gel and afterwards stored at −80 °C. Urediniospores were multiplied on Italian ryegrass genotypes known to be highly susceptible to stem rust. To do so, urediniospores were removed from storage at −80 °C and given a heat shock treatment for five minutes in a 42 °C water bath. A spore suspension, produced by adding spores to 150 ml Dodecan 99 + % (Sigma-Alderich, St. Louis, Missouri, USA; approximate spore density = 1,250,000 urediniospores ml^−1^), was then sprayed on new clones (produced by cloning existing plants two to four weeks prior to inoculation) of the susceptible genotypes. Inoculated plants were left for one hour in darkness at room temperature and were then put under completely closed plastic hoods in a tray filled with water to create high humidity. Plants were placed in a climate chamber (CLIMECAB 1400, Kälte 3000, Landquart, Switzerland) at a humidity of 70%. After 24 hours in darkness at 22 °C, a day-night cycle of 15 h daylight (150 µmol m^−2^ s^−1^ photosynthetically active radiation measured at plant base level) at 22 °C and 9 h dark at 20 °C was applied. Eight to 10 days after inoculation, first stem rust symptoms appeared on the adaxial side of the leaves. Urediniospores were from then on aspirated regularly for twelve weeks with a self-made cyclone spore collector. After collecting, the urediniospores were dried for one week over silica gel and stored at −80 °C.

#### 2.3.2 Leaf segment test (LST)

The LST was performed according to the protocol of Lellbach (1994) with minor modifications as described below. Square petri dishes (120 x 120 x 17 mm, petri dish square, Greiner Bio-One, Kremsmünster, Austria) were filled with 50 ml 5% agar complemented by 30 mg l^−1^ benzimidazol (Sigma-Alderich, St. Louis, Missouri, USA). In total, four leaf segment replicates were tested for each plant. Two technical replicates, derived from the same leaf of the same plant, were placed in the same petri dish. Two biological replicates, derived from two different leaves of the same plant were placed in two different petri dishes. Leaf segments with a length of 28 mm were taken from the youngest mature leaf and placed with the abaxial side of the leaf touching the agar of the petri dish. To compare infection levels between petri dishes, leaf segments of four control genotypes with high, moderate and low susceptibility to stem rust were added to each petri dish. The pre-multiplied urediniospores stored at −80 °C were treated by heat shock for five minutes at 42 °C as described above. For inoculation, petri dishes with leaf segments were placed under a settling tower (size 12.3 cm x 12.3 cm, 40 cm height). Urediniospores were then dispersed within the settling tower using a compressed air duster (0.5 bar) and were allowed to settle down onto the petri dishes for four minutes. The petri dishes were then covered with a lid sprayed with water, wrapped into a wet towel and packed in a black plastic bag. Leaf segments were then incubated for 24 h in the darkness in the climate chamber (CLIMECAB 1400, Kälte 3000, Landquart, Switzerland) at 25 °C. After 24 h, petri dishes were unwrapped and kept for eight days at a day-night cycle of 15.5 h daylight (100 µmol m^−2^ s^−1^ photosynthetically active radiation) at 25 °C, 60% relative humidity and 8.5 h dark at 25 °C. Disease symptoms were scored eight days after inoculation using a scale from 1 to 9, defined as: 1 = no pustules, 2 = 1 - 2 pustules, 3 = 3 - 5 pustules, 4 = > 5 pustules with weak sporulation, 5 = > 15 pustules and some sporulation, 6 = thin equally distributed sporulation, 7 = thin equally distributed sporulation with some strongly infested areas, 8 = leaf segment equally covered with pustules, 9 = leaf segment fully covered with pustules (Beckmann 2010).

### 2.4 Statistical analysis of phenotypic data

Data obtained from the field trials across all three environments (i.e., combinations of location and year) were analyzed using a linear mixed-effect model

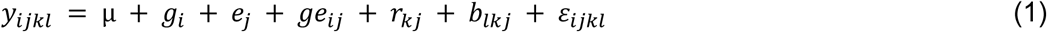

where *y_ijkl_* represents the observation for trait *y* on a single plant basis, *µ* denotes the overall mean, *g_i_* the effect of genotype *i*, *e_j_* the effect of environment *j*, *ge_ij_* the interaction effect of genotype *i* with environment *j*, *r_kj_* the effect of the *k*-th complete block (replicate) nested within the *j*-th environment, *b_lkj_* the effect of the *l*-th incomplete block nested within the *k*-th complete block and *ε_ijkl_* the residual error. To calculate best linear unbiased estimators (BLUEs) for each genotype across all environments, all factors within equation 1 except *b_lkj_* were taken as fixed. For estimation of the respective variance components, *g_i_* and *ge_ij_* were additionally taken as random effects in a second model. For the analysis of data from each environment separately and to calculate BLUEs for each genotype per environment, equation 1 was reduced by the factors *e*_j_ and *ge_ij_*. Broad-sense heritability (*h^2^*) was calculated as

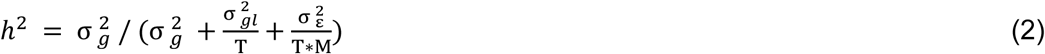

with 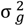 denoting the genotypic variance, 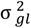 the genotype-by-environment interaction variance and 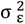 residual variance. T (= 3) represents the number of environments and M (=2) the number of complete blocks (replicates) per environment.

Analysis of the LST data was conducted using the model

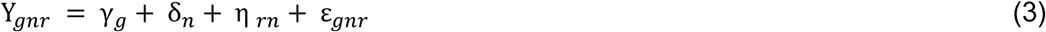

where Y*_gnr_* is the rust score for the *g*-th genotype from the *r*-th replicate nested within the *n*-th petri dish, γ_g_ is the effect of the *g*-th genotype, δ_n_ is the effect of the *n*-th petri dish, η_rn_ is the effect of the r-th replicate within the *n*-th petri dish and ε_gnr_ is the residual error per experimental unit. For calculating the BLUEs for genotypic means from LST data, all factors from equation 3 except η*_rn_* were taken as fixed effects.

For calculation of the variance components, γ*_g_*, δ*_n_* and η*_rn_* were taken as random effects. For the LST, broad-sense heritability (*h^2^*) was calculated as

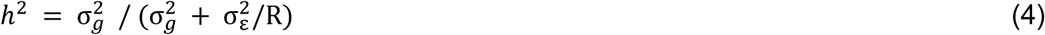

BLUEs were used to calculate the Pearson correlation coefficients among each trial (field and LST) and their significance was tested using a two-tailed t-test.

All analysis were conducted using RStudio (v2025.02.28; (Posit Team 2024) and R (v4.4.3; R Core Team, 2024) and with the packages dplyr (v1.1.4; (Wickham et al. 2023), ggpubr (v0.6.0; (Kassambara 2023), lmerTest (Kuznetsova et al. 2017), lme4 (v1.1.35.5; Bates et al., 2015), emmeans (Lenth 2023), ggplot2 (v3.5.1; (Wickham 2016) and tidyverse (v2.0.0; Wickham et al., 2019).

### 2.5 Genomic analyses

The genotypic matrix of the NAM population was imputed based on whole genomic sequencing data of the NAM founders as described in Kiesbauer et al. (2025) and after filtering it consisted of 3,199,253 SNPs.

#### 2.5.1 Genome wide association analysis

The BLUEs from the field data phenotyping as well as from the LST were separately used for GWAS analysis with GENESIS (v2.24.2; Gogarten et al., 2019). The genetic relationship matrix was calculated and implemented in the mixed linear model (MLM) as a covariance structure for random effects. Based on the low population structure observed (Kiesbauer et al. 2025), no additional covariates were used to control for population structure in the model. To determine significant associations, a Bonferroni-corrected *p*-value of 1.05 x 10^−8^ (α = 5%) was used.

#### 2.5.2 Identification of candidate genes for resistance to stem rust

SnpEff (v4.3; Cingolani et al., 2012) was used to analyse the effect of each marker. All markers above the Bonferroni threshold (α = 5%) having a high or moderate effect in the coding regions of the reference genome (Chen et al. 2025) were inspected in more detail. Based on the linkage disequilibrium (LD) decay of this NAM population (Kiesbauer et al. 2025), the gene annotations located 14.4 kb up- and downstream of these significantly associated markers were evaluated further (Table 3). To find information on the putative function of these genes, the nucleotide sequence of these candidate genes was compared to other crop genomes (*L*. *rigidum*, *L. perenne*, *Oryza sativa*) using BLAST+ (Camacho et al. 2009) in NCBI using the nucleotide database (Sayers et al. 2021). Significantly associated SNPs (Bonferroni threshold with α = 0.05) in coding regions were identified and the corresponding genes were declared as candidate genes and further discussed. SNPs not meeting the Bonferroni threshold, but with a *p*-value lower than *p* = 1.0 × 10^−6^ were described in Supplementary Table 1.

#### 2.5.3 Estimation of haplotypes of candidate gene *Chr7.32208*

To capture more genetic variation within our stem rust resistance candidate gene *Chr7.32208* on chromosome 7, haplotypes were defined. The ddRAD genotypic data of the NAM population and the 23 diverse founders were used to define haplotypes. The data were processed as described in Kiesbauer et al. (2025), but without using the founder whole genome sequencing data to impute the genotypic matrix of the NAM population. The genotypic matrix was filtered to include only genotype calls with a quality score above 30, a minimum read depth of three, and at least 1% of the accessions genotyped per SNP. This resulted in a total of 324,801 SNPs, hereafter reported as unimputed genotypic matrix. To define the haplotypes at the significantly associated SNP to stem rust on chromosome 7 (position 248,216,063 bp), we used SNPs located within the interval of 248,224,751 – 248,225,285 bp. This region contains the predicted gene *Chr7.32208*. Within this defined interval, 17 markers displayed a haplotype. For clustering, the genotype matrix was translated to numeric values. For the 23 diverse founders of the NAM population, the Euclidian distance was calculated and hierarchical clustering according to the Ward.D2 method performed (Ward 1963; Murtagh and Legendre 2014). The resulting number of haplotype clusters among the founders was used to define the number of haplotype groups for the cluster analysis of the NAM F_2_ population.

## Results

### 3.1 High variability for stem rust resistance

The three field trials showed substantial differences in stem rust infections with an average score of 4.3 in environment ZH20, compared to 2.4 and 2.2 in ZH21 and OE21, respectively (Table 1). The distribution of the stem rust scores in ZH21 and OE21 was comparable, being strongly right-skewed with many plants showing low scores (Fig. 1). In contrast, a bimodal distribution was observed for ZH20, with two peaks at low (< 2) and high (> 7) scores. The climatic conditions varied among the different trials, where temperature was higher and rainfall lower in ZH20 (average daily temperature 18.4 °C, total precipitation 94.6 mm, calculation beginning with the flowering of the first plant and ending with the first harvest date within the trial) compared to ZH21 (17.9 °C, 331.0 mm) and OE21 (18.1 °C, 172.2 mm). The LST displayed a range of intermediate to low stem rust scores with values per single leaf sample not exceeding 7. On average, the LST had a score of 2.52 (Table 1). Genotypic variance was highest in ZH20, the environment with the highest rust scores, followed by ZH21 and OE21, and the lowest genotypic variance was observed for the LST (Table 1). Broad-sense heritability for stem rust was generally high, ranging from 0.73 (LST) to 0.89 (ZH20). All tested correlations among environments as well as LST were significant with *p*-values *p* < 0.001 (Table 2). BLUEs of rust scores per genotype were highly correlated among the three field trials (with correlation coefficients ranging from 0.64 to 0.88), while correlation coefficients of BLUEs from the LST with the BLUEs of the individual field trials were low, ranging from 0.24 to 0.42.

**Fig. 1.**
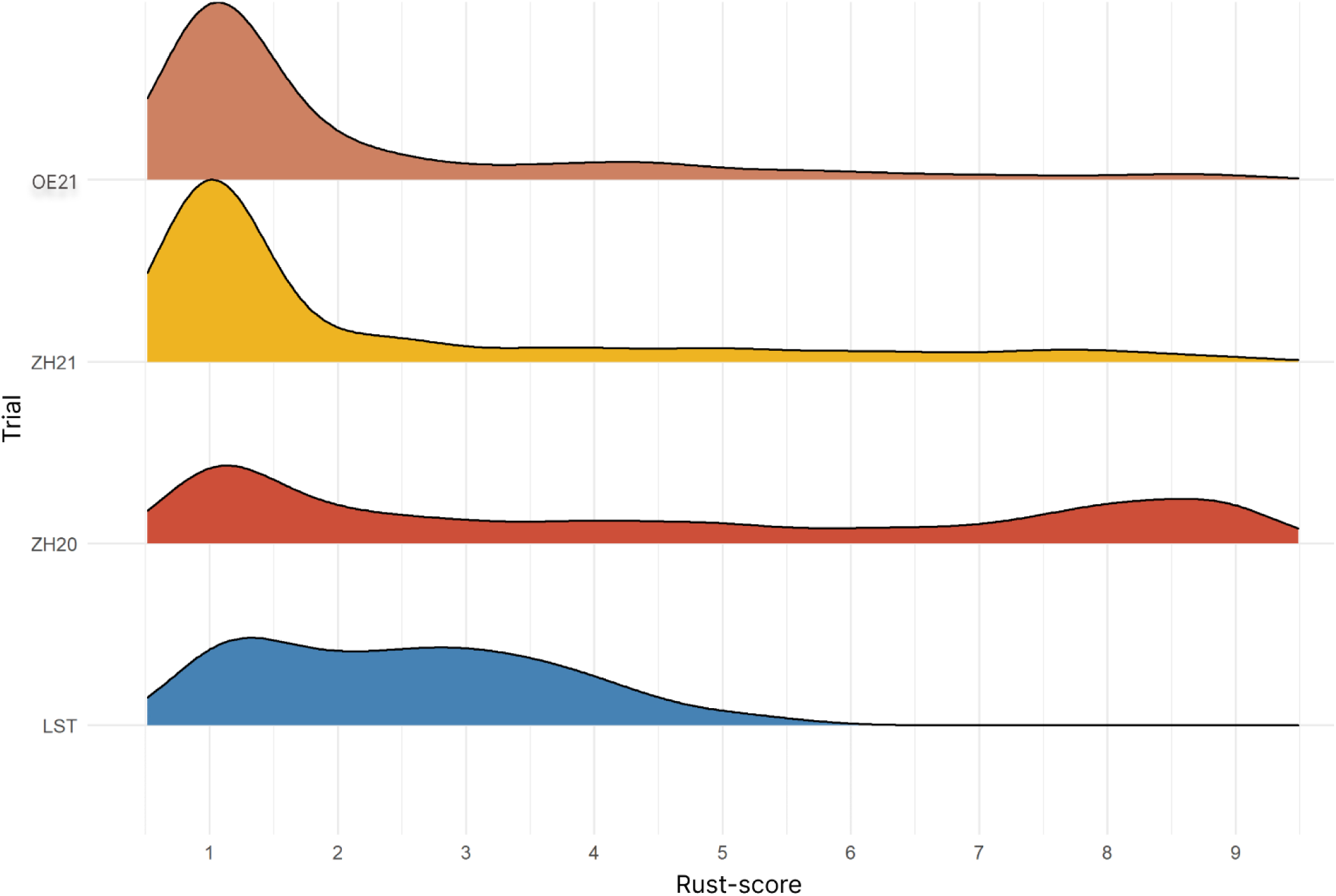
Distribution of stem rust scores in the nested association mapping population for the three field trials (ZH20, ZH21, OE21) and the in vitro leaf segment test (LST) using loess regression for smoothing. Stem rust occurrence in the field trials was scored from 1 (no symptoms) to 9 (more than 75% of the plant is covered with stem rust). Stem rust occurrence for the LST was also scored from 1 (no pustules) to 9 (leaf segment fully covered with pustules).

**Table 1.**
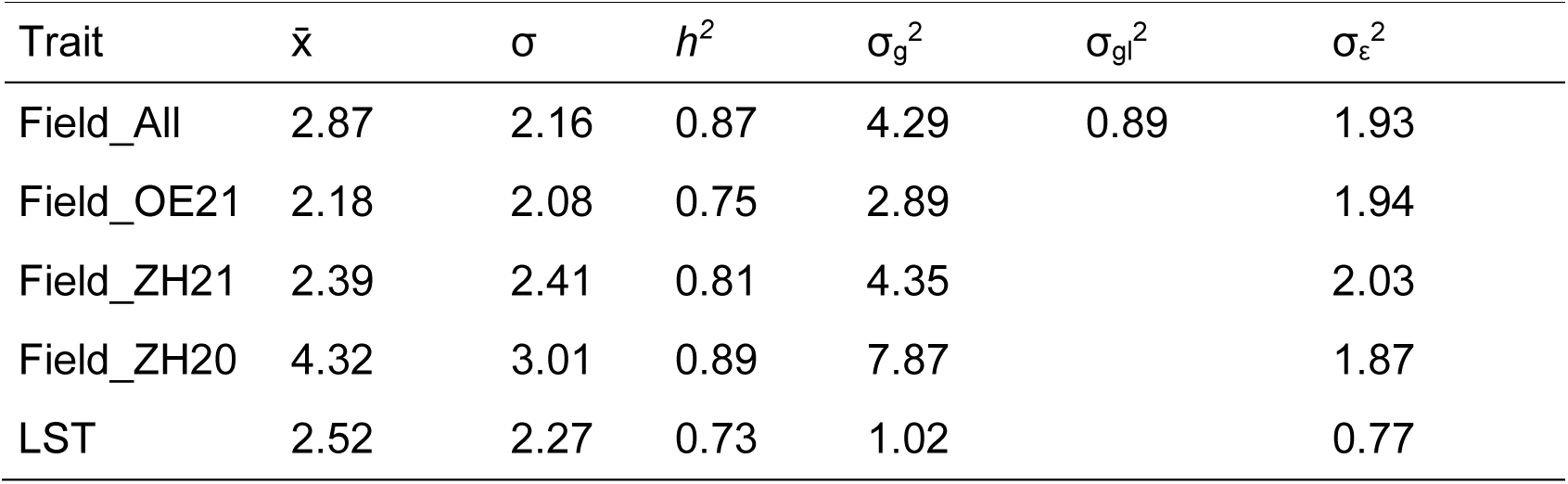
Summary statistics of stem rust resistance in 708 individuals of a nested association mapping population for Italian ryegrass (*Lolium multiflorum* Lam.). Mean (x̄), standard deviation (σ), broad sense heritability (*h^2^)*, genotypic variance (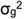), genotype by environment interaction variance (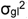) and residual variance (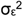) are shown for each environment (Field_ZH20, Field_ZH21 and Field_OE21) and the leaf-segment test (LST) separately and for all field trials together (Field_All).

**Table 2.** Pearson’s correlation coefficients between best linear unbiased estimators of the three field trials Field_ZH20, Field_ZH21 and Field_OE21 (combined and separately). All values are highly significant with *p*-values < 0.001.

| Trait | Field_All | Field_OE21 | Field_ZH21 | Field_ZH20 |
| --- | --- | --- | --- | --- |
| Field_OE21 | 0.83 |  |  |  |
| Field_ZH21 | 0.88 | 0.64 |  |  |
| Field_ZH20 | 0.92 | 0.66 | 0.71 |  |
| LST | 0.42 | 0.24 | 0.32 | 0.43 |

### 2.6 Two main loci associated with stem rust resistance

SNP calling resulted in 3,199,253 SNPs evenly distributed across all seven chromosomes (Supplementary Fig. 1). No significant marker-trait associations were found using LST data (data not shown). The MLM approach using the BLUEs of the combined field data revealed two loci significantly associated with stem rust resistance (Fig. 2). The *λ*–value calculated from the corresponding quantile-quantile plot (i.e., the median of the resulting chi-squared test statistics divided by the expected median of the chi-squared distribution) was 0.97, hence there was no indication for inflation.

**Fig. 2.**
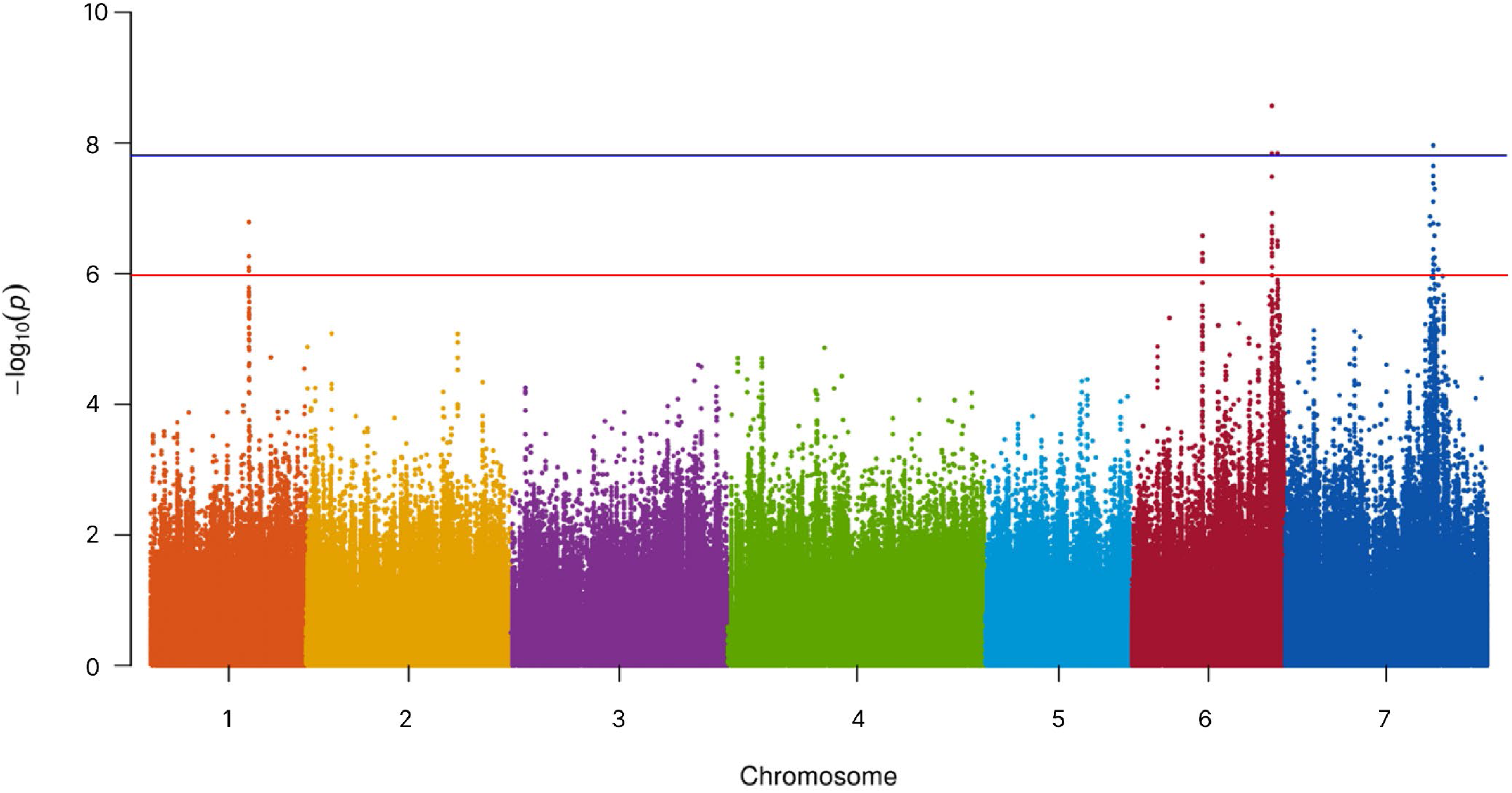
Manhattan plot generated from a nested association mapping population of Italian ryegrass (*Lolium multiflorum* Lam.) consisting of 708 plants and 3,199,253 SNP markers. Stem rust data from three field trials (Field_ALL BLUEs) were analysed using a mixed linear model approach (Segura et al. 2012). The blue line indicates the Bonferroni-corrected significance threshold of 5%. The red line indicates a *p*-value of *p* = 1.0 × 10^−6^.

A total of four SNPs were identified in two main regions above the Bonferroni threshold, three on chromosome 6 and one on chromosome 7 (Fig. 2). Each of the four SNPs significantly associated with resistance were located on a different predicted gene model (Table 3). On chromosome 7, a SNP at position 248,216,063 bp significantly associated with stem rust was found within gene *Chr7.32208.* Of the 13 SNP markers that had the lowest *p*-value on chromosome 7, 12 were found within gene *Chr7.32208.* This gene has a predicted function as a receptor-like serine/threonine kinase (Fig. 3). Further, two of the SNPs within the gene *Chr7.32208* are predicted to result in a missense variation, which led to an amino acid change in the coding sequence. The SNP with the lowest *p-*value on chromosome 7 leading to an intron variation in *Chr7.32208* explained 4.78% of the total phenotypic variance for stem rust and had an effect size of β = −0.88 (Table 3). No other genes were annotated within the LD decay range of 14.4 kb.

**Table 3.**
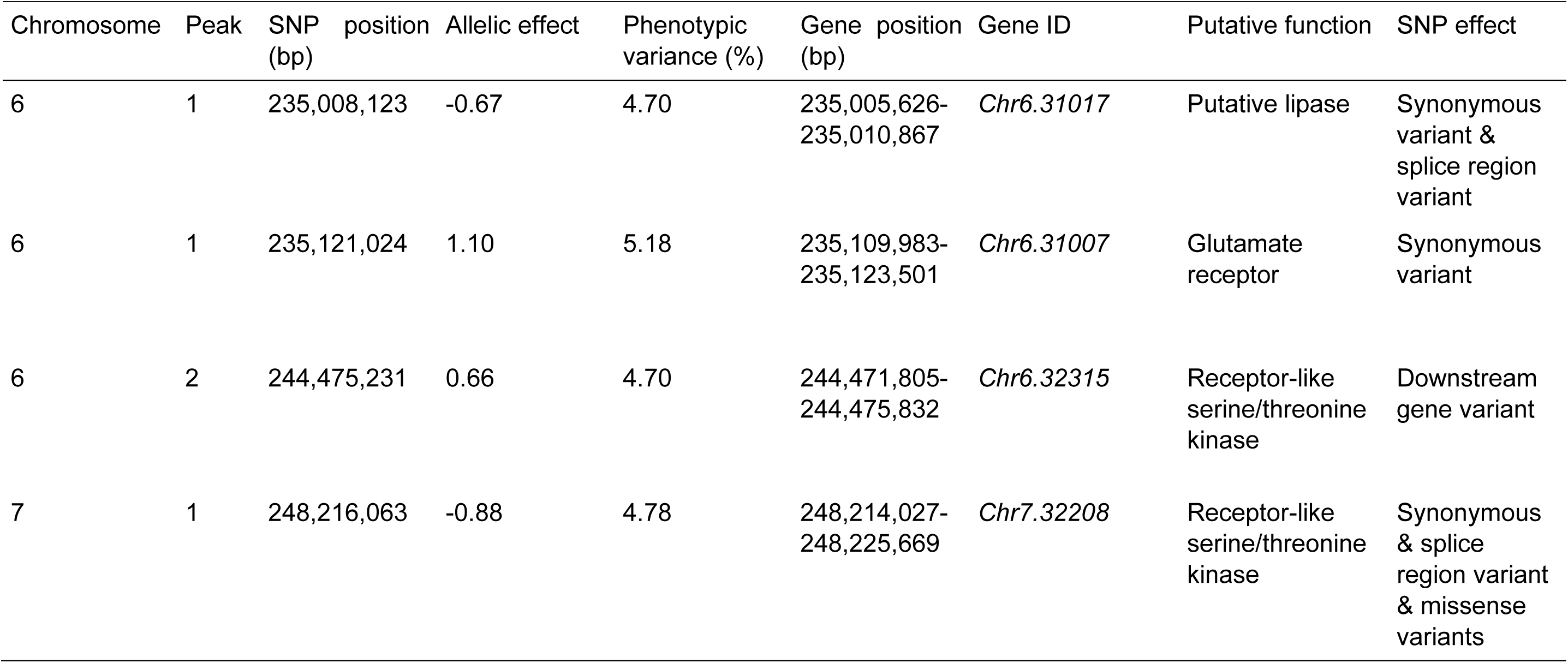
Identification of single nucleotide polymorphisms (SNPs) significantly associated with stem rust resistance (Bonferroni threshold, α = 5%) within predicted genes in the reference genome cv. ‘Rabiosa’ (Chen et al. 2025) in a nested association mapping population of Italian ryegrass (*Lolium multiflorum* Lam.) SNP position indicate the SNP significantly associated with stem rust. Gene name represents the name of the gene in the reference genome. The gene function was revealed by using basic local alignment search tool of NCBI. The allelic effect means the effect of alternative allele.

**Fig. 3.**
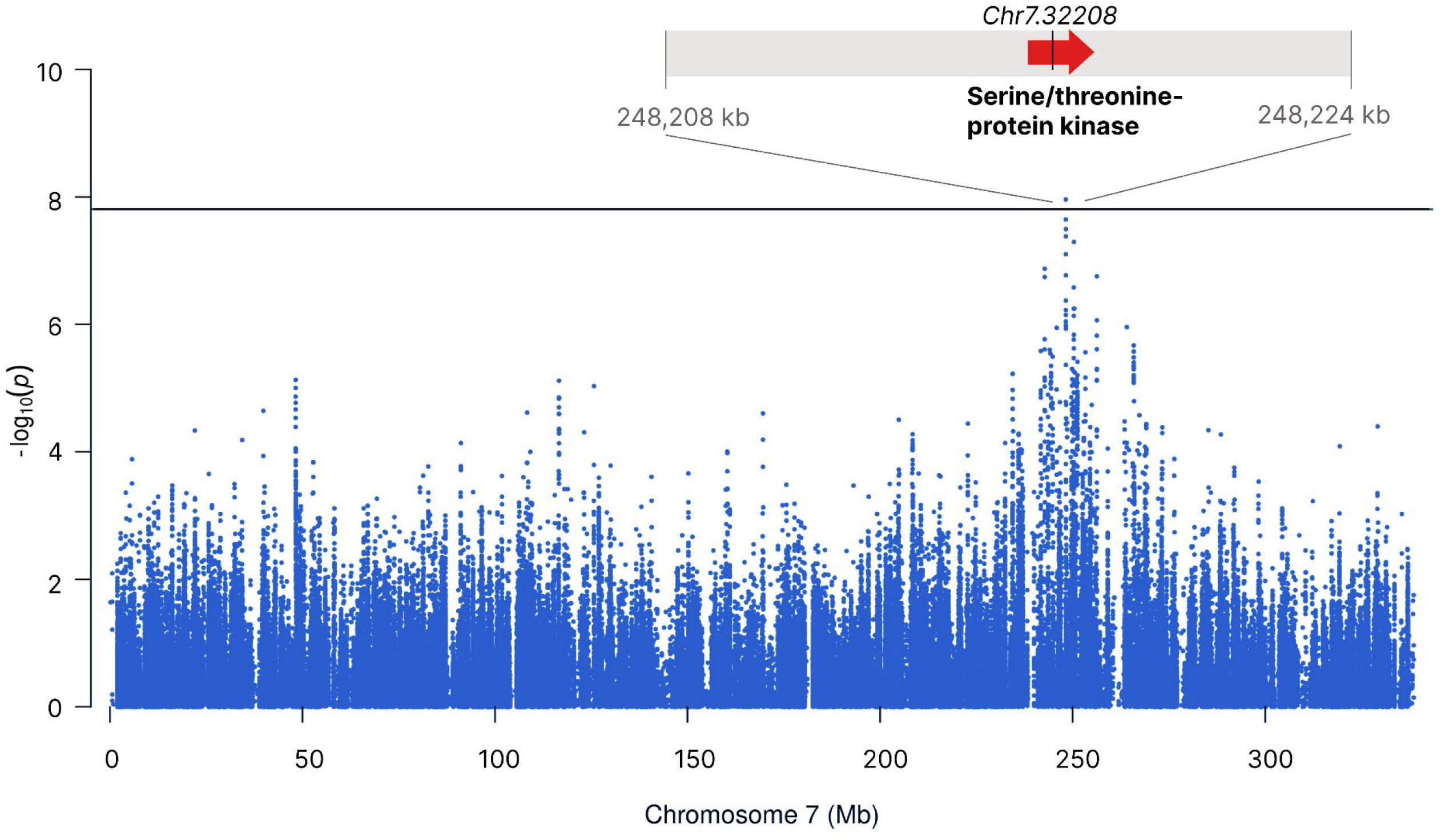
Genome-wide association analysis for stem rust resistance on chromosome 7 using mixed linear model (MLM). The x-axis shows chromosome 7 in Mb whereas the y-axis shows – log10 *p*-values. The black horizontal line indicates the Bonferroni-corrected threshold of 5%. The annotated gene 14.4 kb up- and downstream from the marker with the lowest *p*-value of this peak is shown. The SNP marker significantly associated with stem rust resistance, displayed as a black vertical line, is within a putative receptor-like serine/threonine kinase, which are known to be involved in disease resistance. In total 12 markers within the putative resistance gene showed a very low although not significant *p*-value.

In total, nine different haplotype clusters within the gene *Chr7.32208* were identified based on the ddRAD sequencing data of 23 NAM founders. In the NAM population, haplotype cluster 1 and haplotype cluster 7 had the highest number of haplotypes, 322 haplotypes and 358 haplotypes, respectively. Haplotype cluster 1 and haplotype cluster 8 resulted in a higher resistance to stem rust compared to the other haplotypes (Fig. 4). Most of the NAM parents as well as the diverse founders were highly heterozygous also and, therefore, sorted into two different haplotype clusters. Homozygous plants, which harbor haplotype 1 twice, showed significantly fewer symptoms of stem rust disease than heterozygous plants that harboured haplotype 1 only once. (Supplementary Fig. 3, *p*-value < 0.001). The first peak on chromosome 6 had two significant SNP associations at position 235,008,123 bp and at position 235,121,024 bp, respectively (Table 3). The SNP at position 235,008,123 bp explained 4.70% of the phenotypic variance and had an effect size of β = −0.67 (Table 3). This SNP was found in the predicted protein coding sequence of gene *Chr6.31007*. This gene was predicted to have a function as glutamate receptor (Fig 5). Within a region of 14.4 kb up- and downstream of the SNP significantly associated with the trait, a diacylglycerol kinase gene (*Chr6.31006*) was identified. At position 235,121,024 bp, another gene (*Chr6.3017)* was predicted and the SNP significantly associated with stem rust resistance showed a predicted synonymous variation within the protein coding region. This SNP explained 5.18% of the phenotypic variance and had an effect size of β = 1.10 (Table 3).

**Fig. 4.**
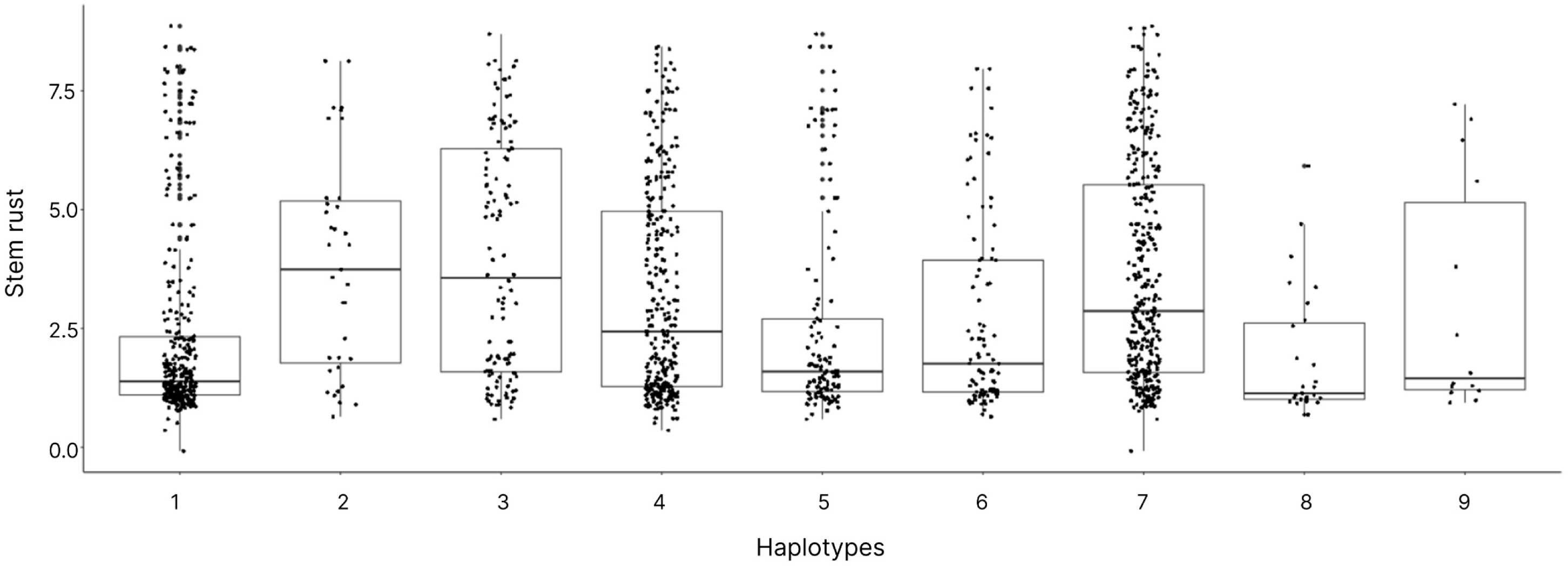
Haplotypes clusters within gene *Chr7.32208* plotted against stem rust scores on the y-axis. The gene has a predicted function as a receptor-like serine/threonine kinase, which is known to be involved in resistance to stem rust. As phenotypic stem rust data, the best linear unbiased estimators from all three field trials were used (Field_All BLUEs). The haplotypes are defined based on 17 SNP markers around the SNP marker with the lowest *p*-value on chromosome 7.The 24 founder genotypes showed nine different haplotypes at this locus.

**Fig. 5.**
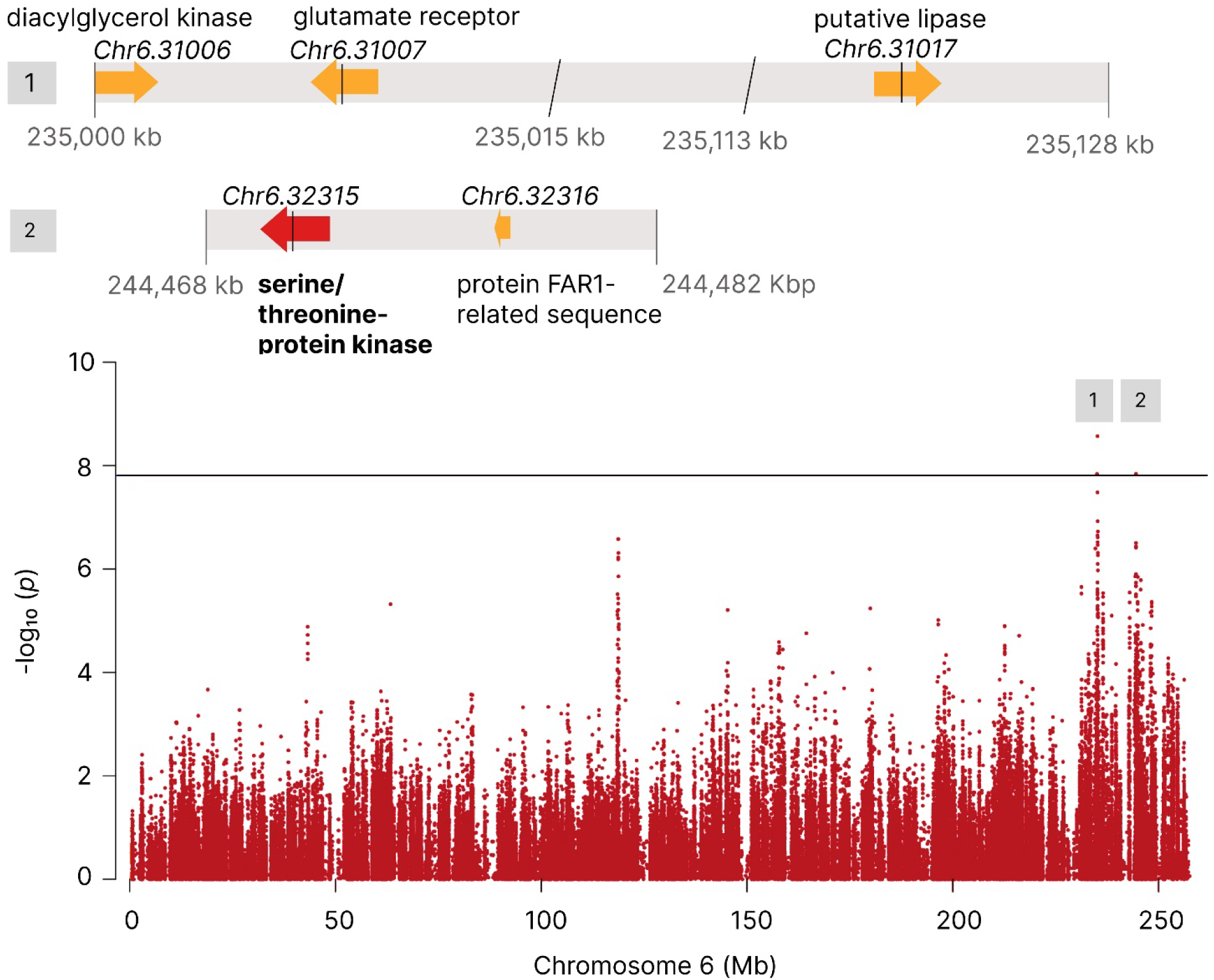
Genome-wide association analysis for stem rust resistance on chromosome 6 using mixed linear model (MLM). The x-axis shows chromosome 6 in Mb whereas the y-axis shows – log10 *p*-values. The black horizontal line indicates the Bonferroni-corrected threshold of 5%. Annotated genes 14.4 kb up- and downstream of these peaks are shown within interval from 235,000 kb - 235,015 kb, from 235,113 kb - 235,128 kb and from 244,468 kb - 244,482 kb, respectively. All SNPs significantly associated with stem rust are indicated by black vertical lines. A putative resistance gene containing a significantly associated SNP is displayed in red. Other genes predicted within the region are shown in yellow.

The second peak on chromosome 6 had one SNP significantly associated with stem rust at position 244,475,231 bp (Fig. 5). The SNP explained 4.70% of the phenotypic variance and had an effect size of β = 0.66 (Table 3). This SNP was found in the predicted intron of gene *Chr6.32315*, having a function as a receptor-like serine/threonine kinase. Within a region of 14.4 kb up- and downstream of the SNP significantly associated with the trait, a gene with a protein FAR1-RELATED SEQUENCE was also predicted (*Chr.32316*).

Further peaks on chromosomes 1 and 6, defined by having a higher *p*-value than *p*= 1.0 × 10^−6^, although not significant associated with stem rust, are described in Supplementary Table 1. Within the 14.4 kb LD window, one gene was annotated on chromosome 6, having an unknown function.

## Discussion

Using a NAM population consisting of 708 plants, we identified two highly promising candidate genes for stem rust resistance, which are predicted to function as serine-threonine protein kinases. As the NAM population was developed using advanced breeding germplasm with large phenotypic and genetic diversity, the loci associated with stem rust resistance may be directly used in the breeding germplasm.

In all three different environments, severe infections with natural stem rust occurred, whereby ZH20 environment showed the highest infection scores. The infection pressure of stem rust in forages can be explained by the weather conditions, where temperature and moisture play a key role (Pfender 2009). Most probably the high disease pressure in ZH20 can be explained by a higher daily average temperature compared to ZH21 and OE21. Rust scoring in the field is time-consuming, and the occurrence of stem rust depends on optimal weather conditions. On the other hand, artificial inoculation of leaf segments with rust pathogens is described as an easy and fast method for phenotyping resistance to fungal diseases like crown rust in perennial ryegrass (Studer et al. 2007; Schubiger and Boller 2016). In comparison to field screenings, artificial inoculation using an LST has the advantage that the phenotyping can be done independently of the field season with single spore isolates or defined spore mixtures from pathovars collected all over the world. However, our results showed a low correlation between field and LST and therefore LST is less suitable for stem rust resistance mapping and cannot replace scoring in the field trials. For the LST in this study, a mixture of spores was collected from the field trial in 2020. Therefore, differences between stem rust populations used in the field trials and in the LST are unlikely. The phenotypic variation observed with the LST was lower when compared to the phenotypic variation of stem rust present in our field trials. In LST the positive controls, as well, showed fewer symptoms compared to the positive controls in the field trials. One reason could be that the used inoculation protocol was originally described for crown rust inoculation and, therefore, the climate chamber conditions were not optimal adapted for stem rust. Also, the correlation between all field scoring data and the LST scoring data was rather low (*r* = 0.42). One explanation could be that under natural conditions (e.g. field scorings) the main disease symptoms of stem rust in the field occur on leaf sheaths and culms which could be different to observing leaf symptoms by using LST. Another inoculation method, which can be carried out under controlled conditions, was applied to whole plants under greenhouse conditions (Pfender et al. 2011). However, this method is very laborious and was so far not compared with field data.

In comparison to the LST association mapping, where no markers significantly associated with stem rust resistance were found, our field data revealed four SNPs above the Bonferroni threshold associated with stem rust resistance. Three of them mapped on chromosome 6 and one on chromosome 7 (Fig. 2). No common markers were used within our study compared with previous QTL mapping studies and, therefore, a comparison between this and other findings of markers associated with stem rust resistance cannot be done. An in silico BLAST with the published co-segregating marker of the locus *LpPg1* (Bojahr et al. 2016) revealed a hit on chromosome 7 on position 285,530,841 bp using reference genome cv. ‘Rabiosa’ (Chen et al. 2025), 37 Mb downstream of the region that showed significantly associated markers in our experiment. No gene is predicted within this position. Also, the closest markers from pathotype-specific resistance locus on LG7 published by Pfender and Slabaugh (2013), is more than 22 Mb away from our SNP significantly associated with stem rust on chromosome 7 (data not shown). Based on the large distance to the previously published QTLs found for stem rust, we can assume that we have found novel loci of resistance.

Until now, studies on stem rust in ryegrass only suggest markers which are co-segregating with a stem rust resistance locus. However, in case of recombination between the marker used for selection and the causative resistance gene, the linkage can be lost and would results in false positive selection (Andersen and Lübberstedt 2003). For example, Beckmann (2010) showed two flanking markers to the resistance locus *LpPG1* with a distance of 2.6 cM and 6.7 cM, respectively. After several breeding cycles, the co-segregation of resistance and a single marker is expected to be lost at least in several plants. By defining haplotypes within the receptor-like serine/threonine kinase candidate gene *Chr7.32208*, the spectrum of genetic variation within the candidate gene can be increased and, therefore, the resolution for differentiating resistant and susceptible genotypes can be increased (Qian et al. 2017; Hamazaki and Iwata 2020). Haplotypes from cluster 1 and 8 are superior resulting in the highest stem rust resistance compared to the other clusters. By screening for these two superior haplotypes within a breeding population, the risk of selecting plants carrying the marker but not the desired resistance allele is reduced. Yet, the development of functional markers requires precise knowledge of the causal resistance origin.

Here, four candidate genes were identified, three on chromosome 6 and one on chromosome 7, showing significant associations with resistance to the field inoculum of stem rust across three environments (Table 3). Among the four candidate genes, two belong to the receptor-like serine/threonine protein kinase family (Table 2). Receptor-like serine/threonine kinase and nucleotide-binding site leucine rich repeat NBS-LRR belong both to the group of receptor protein kinases. NBS-LRR genes are well known to be involved in plant disease resistance (McHale et al. 2006). NBS-LRR have been previously identified to confer resistance to stem rust in *L. perenne* (Bojahr et al. 2016). Receptor-like serine/threonine kinases are also reported to be involved in fungal pathogen resistance (Cao et al. 2011). For example in barley, *Rpg1* confers resistance to several, but not all, pathotypes of stem rust caused by *P. graminis* f.sp. *tritici* and belongs to the receptor-like serine/threonine kinase (Brueggeman et al. 2006). There, *Rpg1* resistance is dominant and reported to be durable in barley (Brueggeman et al. 2002). The fact that a receptor-like serine/threonine kinase is involved controlling resistance to stem rust in another crop supports our hypothesis that our receptor-like serine/threonine kinase resistance candidate genes could be involved in resistance against stem rust in Italian ryegrass. Moreover, the SNP markers within these two candidate genes resulted in downstream gene variant or a missense variant (Table 3). Both variants may influence the expression of the gene. However, regulatory elements or transposable elements for example could play a role in resistance. To clarify whether and which of these two receptor-like serine/threonine kinase genes are involved in stem rust resistance, further investigations are needed. Gene expression (Li et al. 2019; Adolfi et al. 2019) and knockout (Hoffie et al. 2021; Lu et al. 2022; Távora et al. 2022) studies would be an option for functional validation of the candidate genes.

The results of the SNP analysis as well as the haplotype clustering analysis showed that two alleles of the receptor-like serine/threonine kinase resistance candidate gene *Ch7.32208* seem to be linked to an incomplete resistance to the spore mixture in the field. The incomplete resistance can be explained by the mixture of spores involved in natural infection, where most probably more than one pathotype of stem rust is involved. The incomplete resistance could either be due to quantitative resistance to all pathotypes or by qualitative effects, which are limited to some pathotypes within the inoculum. A previous study involving a spore mixture in field experiments produced similar results, with only a few plants displaying complete resistance to natural field infections (Pfender et al. 2011). On the other hand, some single spore isolate inoculations indicated a pathotype specific reaction, which means race specific resistance to specific isolates (Pfender and Slabaugh 2013). To clarify the interaction between fungal pathogens and the plant defense mechanism, further experiments are needed. For example, reinoculation of the same NAM population with one stem rust pathotype would help to better understand the resistance mechanisms involved. However, a mix of spores reflect the reality, and the superior haplotypes can be directly used in a haplotype-based breeding approach.

In conclusion, our study showed that a NAM population with high phenotypic variation of stem rust symptoms due to natural inoculation is a valuable source to find novel resistance sources. In total, four loci were identified, and the SNP markers significantly associated with stem rust were found to be located directly within four annotated genes. Two of these genes are receptor-like serine/threonine kinases, which are known to be involved in stem rust resistance. The superior haplotypes within the receptor-like serine/threonine kinase gene *Chr7.32208* linked to a higher stem rust resistant plant can be directly applied to the breeding program. To study host pathogen interactions as well as to validate the resistance candidate genes, further studies are recommended.

## Data availability

The mapped WGS and ddRAD files are available in the NCBI Sequence Read Archive as BioProject accession number PRJNA1167222.

## Acknowledgements

Genotyping data of the NAM F_2_ population were generated in collaboration with the Genetic Diversity Centre (GDC), ETH Zurich. We thank the Functional Genomic Centre Zurich (FGCZ) for sequencing the NAM founders. This project was supported by the Breeding Foundation DSP-BLW.

## Supplementary files

**Supplementary Fig. 1.**
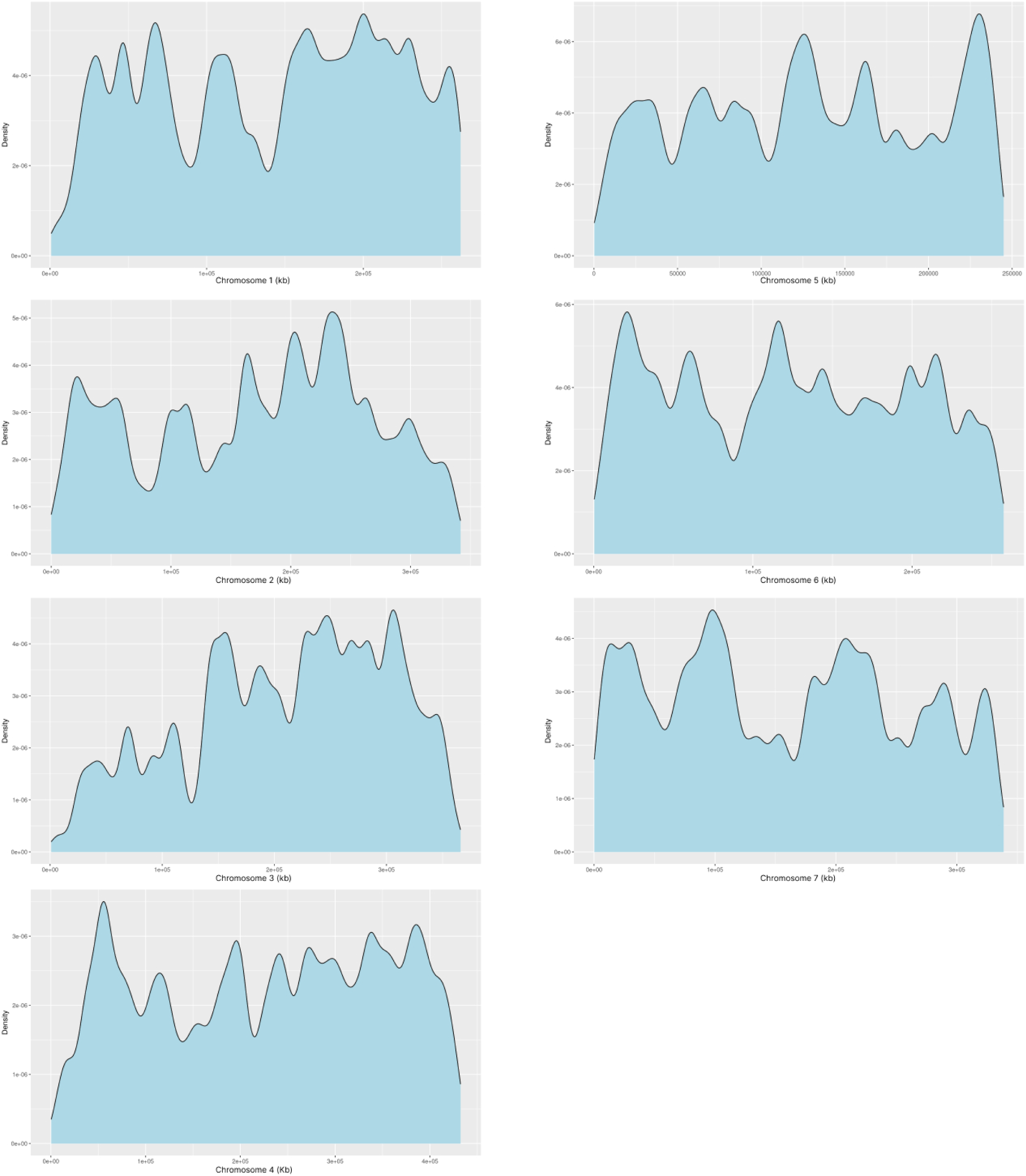
Density of single nucleotide polymorphism (SNP) markers along the seven chromosomes of reference genome cv. ‘Rabiosa’ (Chen et al. 2025). Marker density is shown for each chromosome separately. The x-axes show the length of each chromosome in kb and the y axes display the density of the markers.

**Supplementary Table 1:** Identification of single nucleotide polymorphisms (SNPs) associated with stem rust resistance that have a *p*-value smaller than *p* = 1.0 × 10^−6^ but are below the Bonferroni significance threshold of 5% in a nested association mapping population (NAM) of Italian ryegrass (*Lolium multiflorum* Lam.). The NAM population consists of 708 F2 individuals derived from 24 founder plants exhibiting high variation in stem rust resistance. As phenotypic data, the best unbiased linear estimators from three different environments were used. The genotypic matrix consists of 3,199,253 SNP markers. Gene name represents the name of the gene in the reference genome cv. ‘Rabiosa’ (Chen et al. 2025). The gene function was revealed by basic local alignment search tool. The allelic effect means the effect of alternative allele.

| Chromosome | SNP position (bp) | Allelic effect | Phenotypic variance (%) | Gene position (bp) | Gene name | SNP effect |
| --- | --- | --- | --- | --- | --- | --- |
| 1 | 165,319,368 | 1.26 | 4.00 | - | - |  |
| 6 | 118,700,714 | 1.76 | 3.90 | 118696039-118704239 | *Chr6.15818* | Synonymous variant, intron variant |

**Supplementary Fig. 3.**
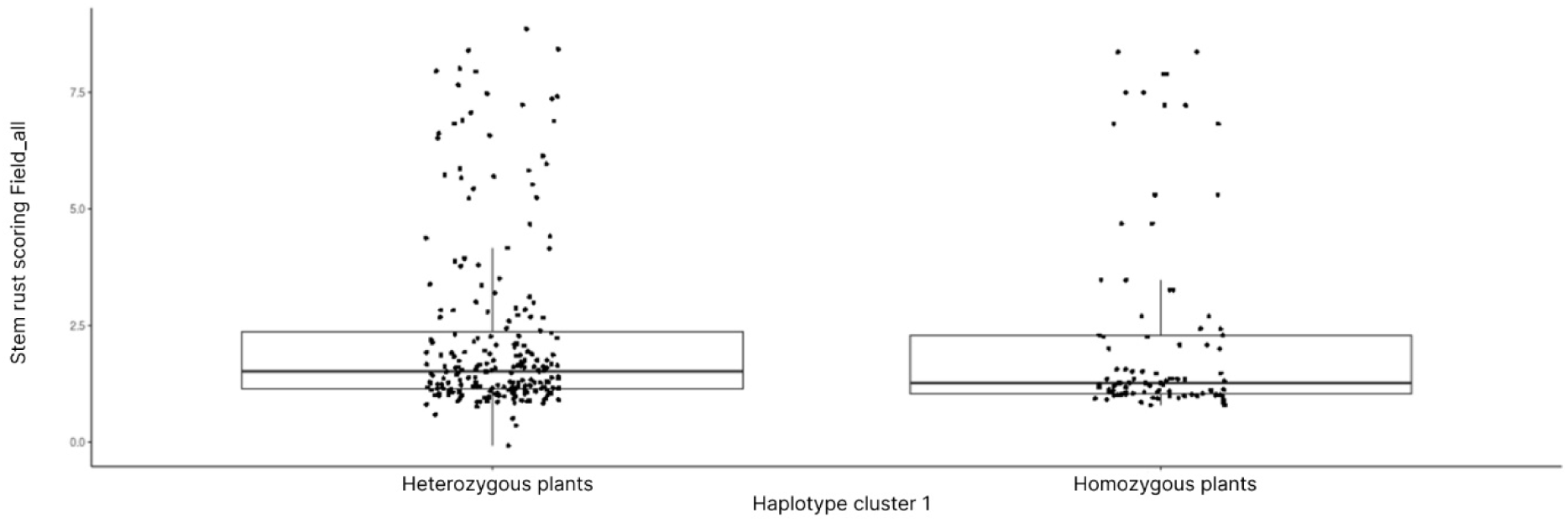
Haplotype cluster 1 divided into two groups. If a plant has two copies of haplotype cluster1, it is considered homozygous. If the plant has only one allele of haplotype cluster 1 and the second one belongs to one of the other haplotype clusters, it is labeled heterozygous plant. The y-axis displays the stem rust scoring for all field data using the best linear unbiased estimator. Homozygous plants showed significantly lower stem rust symptoms than heterozygous plants (*p*-value: < 2.2e-16).

